# Isolation, Identification and Antibiogram Assay of *Escherichia coli* from the Environment of Live Bird Markets in Bangladesh

**DOI:** 10.64898/2026.08.09.743748

**Authors:** Mst. Nahida Akter, Md. Rimon Bhuiyan, Md. Sohel Rana, Rubina Khatun, Anna Purnna Ray, K.M. Mozaffor Hossain

**Author notes:** **Correspondence to: Dr. K.M. Mozaffor Hossain,** Professor, Department of Veterinary & Animal Sciences, Faculty of Veterinary & Animal Sciences, University of Rajshahi, Rajshahi-6205, Bangladesh.

## Abstract

**Background:** Live bird markets (LBMs) may facilitate the persistence and dissemination of Escherichia coli and antimicrobial-resistant bacteria because of intensive bird handling, environmental contamination and inadequate sanitation. However, information on E. coli contamination and antimicrobial susceptibility in LBM environments in Rajshahi District, Bangladesh, remains limited.

**Objective:** This study aimed to determine the prevalence, identify the cultural and biochemical characteristics, and assess the antimicrobial susceptibility pattern of E. coli isolated from water, soil and bird-dropping samples collected from LBMs in Rajshahi District.

**Methods:** A total of 60 environmental samples, comprising 20 water, 20 soil and 20 bird-dropping samples, were collected from LBMs across all ten upazillas of Rajshahi District between January and June 2023. E. coli was isolated and identified using cultural characteristics, Gram staining and biochemical tests. Antimicrobial susceptibility was determined by the Kirby– Bauer disc diffusion method against seven antimicrobial agents using CLSI interpretive criteria.

**Results:** E. coli was detected in 33 of 60 samples, giving an overall prevalence of 55.00%. Prevalence was highest in bird-dropping samples (75.00%), followed by water (55.00%) and soil (35.00%). Among the 33 isolates, resistance was highest to oxytetracycline (78.79%) and amoxicillin (63.64%), followed by ciprofloxacin (48.48%), doxycycline (33.33%), levofloxacin (9.09%), erythromycin (9.09%) and neomycin (6.06%). Sensitivity was highest to neomycin (60.61%), followed by levofloxacin and erythromycin (51.51% each).

**Conclusion:** The high prevalence of E. coli and substantial resistance to several commonly used antimicrobials indicate considerable microbiological and antimicrobial-resistance concerns in LBM environments. Improved sanitation, biosecurity, hygienic poultry handling and prudent antimicrobial use are warranted to reduce environmental contamination and potential transmission of resistant bacteria.

## 1. INTRODUCTION

The poultry industry occupies a central position in Bangladesh’s agricultural economy, serving as a major source of nutrient-dense protein and contributing significantly to rural livelihoods and poverty alleviation. The sector, comprising layer and broiler production together with numerous backyard flocks, has expanded rapidly, with the national poultry population growing at an estimated 0.75% per year between 1970 and 1980 and at nearly 5% per year between 1990 and 2005. Bangladesh currently has an estimated 90,000 licensed poultry farms, although many more remain unregistered, and the Bangladesh Poultry Industries Central Council reports 216 registered parent-stock enterprises supplying roughly 1.48 crore day-old chicks weekly. Poultry meat presently accounts for 22-27% of the country’s animal protein supply and about 37% of total meat production, with average per-capita availability of approximately 136.18 g of meat per day (DLS, 2021).

*Escherichia coli* is a Gram-negative, rod-shaped commensal organism that normally inhabits the lower intestinal tract of humans, poultry and other warm-blooded animals. While the majority of strains are harmless, opportunistic and pathogenic serotypes comprising roughly 10-15% of intestinal coliforms can cause a wide spectrum of disease in poultry and in immunocompromised hosts, including septicaemia, meningitis, endocarditis, urinary tract infection and epidemic diarrhoea (Daini et al., 2005), as well as yolk-sac infection, omphalitis, cellulitis, swollen head syndrome, coligranuloma and colibacillosis (Baergen, 2013). Avian pathogenic *E. coli* (APEC), a member of the extra-intestinal pathogenic *E. coli* (ExPEC) group, typically affects broilers between four and six weeks of age and is characterised by acute fatal septicaemia or subacute fibrinous pericarditis, airsacculitis, salpingitis and peritonitis (Alexander et al., 2017). Certain food-borne serotypes, notably O157:H7, have caused notable outbreaks of gastrointestinal illness over the past two decades (Armstrong et al., 1996).

Beyond its role as a pathogen, *E. coli* is widely used as an indicator organism of faecal contamination in food, water and the wider environment (Akond et al., 2009). Certain populations of environmentally adapted *E. coli* are capable of persisting and multiplying outside the host for extended periods (Jang et al., 2017), and such strains cannot always be distinguished from freshly excreted faecal strains using conventional water-quality testing, so their presence may not necessarily indicate recent faecal contamination. Manure and other animal wastes, slaughterhouse effluent and wastewater treatment discharge are all recognised sources of environmental contamination by pathogenic *E. coli* (Balière et al., 2015). Although extensive research has addressed the clinical features, pathophysiology and diagnosis of pathogenic *E. coli*, comparatively little is known about the ecology and abundance of these organisms once they enter extra-intestinal environments (Kaper et al., 2004), a gap that is of particular concern when the persisting strains carry genes for virulence or antimicrobial resistance.

Antimicrobial resistance (AMR) has become one of the most pressing global public health concerns (Ferri et al., 2017). Antibiotics are extensively used in Bangladesh’s poultry sector both to control infectious disease and as growth promoters (Hassan et al., 2014), and the resulting selection pressure has been strongly implicated in the emergence of antibiotic-resistant *E. coli* (Islam et al., 2018). Using an international database of antibiotic sales, Klein et al. (2018) reported that global antibiotic consumption rose by 39% between 2000 and 2015, from 11.3 to 15.7 defined daily doses per 1,000 inhabitants per day. Selective antibiotic pressure arising from the continual interaction between antimicrobials and bacterial populations is considered one of the principal drivers of the current AMR crisis (Kolar et al., 2001), and inappropriate prescription together with outright overuse of antimicrobials remain the two leading contributors to the problem (Read and Woods, 2014; WHO, 2020).

Because the intestines of warm-blooded animals expose resident *E. coli* to repeated antimicrobial pressure, the organism is subject to strong selection for resistance to the drugs consumed by its host (Looft and Allen, 2012), a pattern that has even been proposed, though ultimately with limited practical value, as a potential tool for host-source attribution of isolates (Harwood et al., 2014). Resistance phenotypes also vary systematically among *E. coli* phylogenetic groups independent of the acquisition of new resistance determinants, indicating that the organism’s genetic background itself shapes its resistance pattern (Tenaillon et al., 2010). Resistant *E. coli* can additionally disseminate resistance determinants to other bacterial species through horizontal gene transfer, acting as a reservoir of transferable antibiotic resistance within the intestinal and environmental microbial community (Salyers et al., 2004). Antimicrobial therapy is widely regarded as the single most important driver of the emergence, selection and spread of antibiotic-resistant organisms in both human and veterinary medicine (Neu, 1992), and the resulting proliferation of resistant strains has been accompanied by increasingly common plasmid-mediated transfer of resistance across bacterial genera (Davies, 1994). Multidrug-resistant *E. coli* is now commonly recovered from both human and animal sources worldwide (Amara et al., 1995), and the large reservoir of drug-resistant, non-pathogenic *E. coli* within the intestine is considered a significant source of resistance genes (Osterblad et al., 2000). *E. coli* has previously been isolated, identified and characterised in Bangladesh from water, milk, eggs, chicken carcasses and calf diarrhoea samples, among other sources (Hasina, 2006).

Live bird markets (LBMs), where consumers purchase live or freshly slaughtered poultry, represent one of the most important nodes of the poultry value chain across South and South-East Asia. Because birds from multiple sources are continually introduced and held together under crowded conditions, and because customers have direct contact with live or freshly processed birds, LBMs create conditions favourable for the persistence and spread of environmental *E. coli* and provide a route through which antibiotic-resistant organisms may enter the human food chain. Despite this, awareness of the potential for LBMs to harbour and disseminate resistant bacteria remains limited in Bangladesh. The present study was therefore designed to determine the prevalence and antibiogram profile of *E. coli* in the water, soil and bird-dropping environment of live bird markets across Rajshahi District, with the following specific objectives:

1. To study the prevalence of *E. coli* in live bird markets of Rajshahi District;
2. To isolate and identify *E. coli* from the live bird market environment of Rajshahi District;
3. To determine the antibiotic sensitivity pattern of each isolate; and
4. To evaluate the public health significance of *E. coli* recovered from live bird markets of Rajshahi District.

## 2. MATERIALS AND METHODS

### 2.1 Study Area and Period

The research was conducted in the Microbiology Laboratory, Department of Veterinary and Animal Sciences, University of Rajshahi, during the period from January to June 2023. Sample collection was carried out at randomly selected live bird markets distributed across all ten upazillas of Rajshahi District, namely Rajshahi Sadar, Puthia, Paba, Tanore, Mohonpur, Durgapur, Baghmara, Godagari, Charghat and Bagha.

### 2.2 Sample Collection

A total of 60 environmental samples were collected aseptically from the live bird market environment for the isolation and identification of *E. coli*, comprising 20 water samples (surface and environment water), 20 soil samples (dust and ground soil) and 20 bird-dropping samples, with 6 samples (2 of each type) collected from each of the ten upazillas. Sterile plastic bags, Eppendorf tubes and cotton swabs were used for sample collection. Sterile cotton swabs, moistened with sterilised normal saline, were used to swab the sample material, immersed in 10 ml of normal saline and subsequently transferred to 60 ml of buffered peptone water, followed by incubation at 37°C for 18 hours. All samples were transported to the Microbiology Laboratory under a maintained cool chain for immediate processing.

**Fig. 1.**
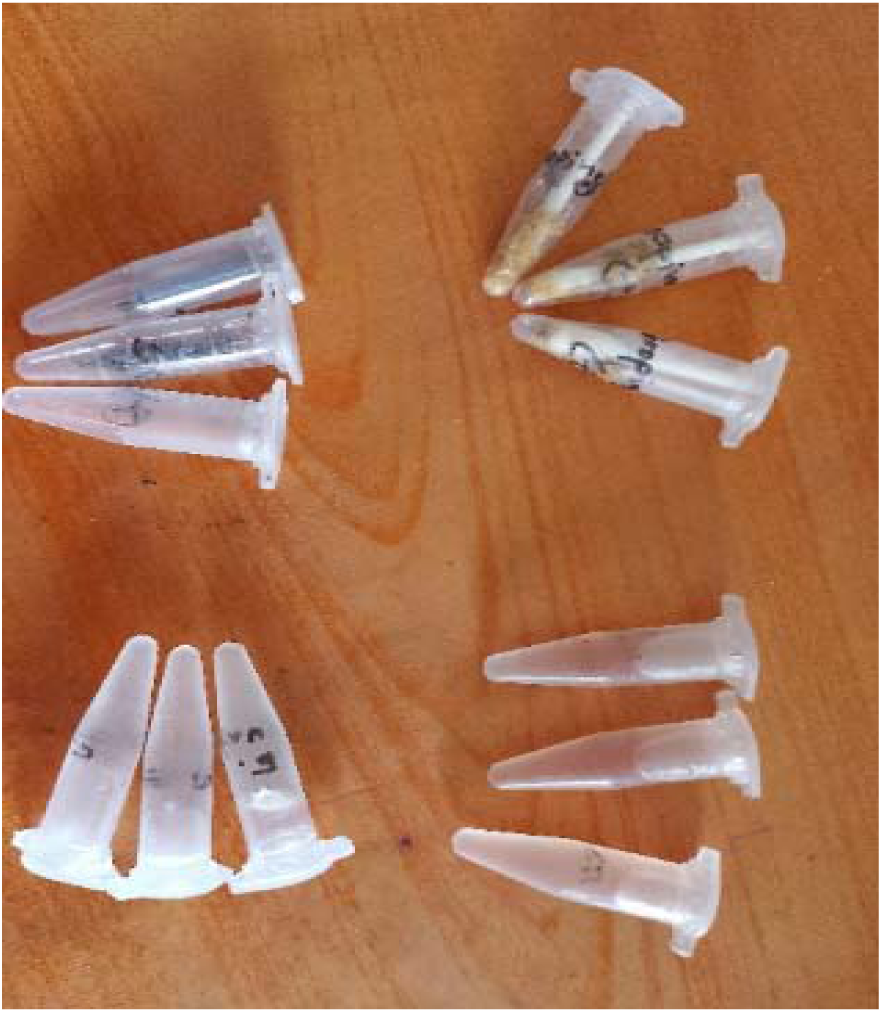
Collection of water, soil and bird-dropping samples from the live bird market environment.

**Table 1.** Distribution of environmental samples collected from live bird markets across the upazillas of Rajshahi District.

| Upazilla | Water | Soil | Bird droppings | Total samples |
| --- | --- | --- | --- | --- |
| Rajshahi Sadar | 2 | 2 | 2 | 6 |
| Puthia | 2 | 2 | 2 | 6 |
| Paba | 2 | 2 | 2 | 6 |
| Tanore | 2 | 2 | 2 | 6 |
| Mohonpur | 2 | 2 | 2 | 6 |
| Durgapur | 2 | 2 | 2 | 6 |
| Baghmara | 2 | 2 | 2 | 6 |
| Godagari | 2 | 2 | 2 | 6 |
| Charghat | 2 | 2 | 2 | 6 |
| Bagha | 2 | 2 | 2 | 6 |
| Total | 20 | 20 | 20 | 60 |

### 2.3 Media, Chemicals and Reagents

Liquid media used in the study included peptone water broth, Methyl-Red and Voges-Proskauer (MR-VP) broth and nutrient broth (NB), while solid media comprised nutrient agar (NA), blood agar (BA), MacConkey agar (MAC), Eosin Methylene Blue (EMB) agar, Xylose Lysine Deoxycholate (XLD) agar, Brilliant Green agar (BGA), Triple Sugar Iron (TSI) agar, Simmons citrate agar (SCA) and Mueller-Hinton agar (MHA). All media were prepared following the manufacturers’ recommended protocols, dispensed into sterilised glassware and sterilised by autoclaving at 121°C and 15 lb/in² pressure for 15 minutes. Standard glassware and equipment, including a hot-air oven, autoclave, incubator, refrigerator, compound microscope, inoculating loops, Petri plates, Durham’s tubes, pipettes and a Bunsen burner, were used throughout the investigation. Isolated bacterial cultures were preserved in 50% sterile buffered glycerin at -20°C, a method that allows organisms to be maintained without alteration of their original characteristics for six months to one year.

### 2.4 Isolation of Bacteria

Following enrichment in nutrient broth, a loopful of the enriched culture was streaked onto EMB agar for the isolation of *E. coli*, and inoculated plates were incubated aerobically at 37°C for 18-24 hours. Presumptive *E. coli* colonies, dark and possessing a characteristic metallic green sheen, were subcultured onto fresh EMB plates to obtain pure cultures, which were incubated for a further 24 hours at 37°C prior to identification.

### 2.5 Identification of Bacteria

#### 2.5.1 Colony Morphology

After 24 hours of incubation, colony morphology, including size, shape, texture, elevation, edge, colour and opacity, was examined and recorded on nutrient agar, EMB agar, MacConkey agar, Brilliant Green agar and XLD agar.

#### 2.5.2 Gram’s Staining

Gram’s staining was performed following the procedure of Cheesbrough (1985). Smears prepared from pure colonies were heat-fixed, stained with crystal violet for two minutes, treated with Gram’s iodine as a mordant for one minute, decolourised with acetone-alcohol, counterstained with safranin for two minutes, and examined under a compound microscope using a 100x oil-immersion objective.

#### 2.5.3 Motility Test

Motility was assessed using the hanging-drop technique. A drop of broth culture was placed on a cover slip and inverted over the concave depression of a hanging-drop slide sealed with Vaseline, and bacterial movement was examined under a 100x oil-immersion objective to distinguish motile from non-motile organisms.

#### 2.5.4 Biochemical Tests

Isolates were characterised biochemically using the catalase test, indole test, methyl-red (MR) test, Voges-Proskauer (VP) test, Triple Sugar Iron (TSI) agar slant reaction, Simmons citrate utilisation test and sugar fermentation reactions with five basic sugars (dextrose, maltose, lactose, sucrose and mannitol), following standard procedures (Cheesbrough, 1985). In the catalase test, a colony was mixed with a drop of 3% hydrogen peroxide and observed for bubble formation. For the indole test, cultures grown in peptone water for 48 hours were treated with Kovac’s reagent and examined for development of a red colour in the reagent layer. The MR test was read following overnight incubation in glucose phosphate broth, a distinct red colour indicating a positive reaction. In the VP test, cultures incubated in glucose phosphate peptone water were treated with creatine and sodium hydroxide, a pink colouration indicating a positive result. Sugar fermentation was assessed by inoculating each sugar medium containing inverted Durham’s tubes and incubating for 24 hours at 37°C, with a colour change and gas production indicating a positive reaction. TSI agar slant reactions were read for slant and butt colour change, gas production and hydrogen sulphide production following stab inoculation and incubation.

### 2.6 Antibiogram Assay (Antibiotic Susceptibility Testing)

The susceptibility of the isolated *E. coli* to seven commercially available antibiotic discs (Hi-Media, India) was determined by the Kirby-Bauer disc diffusion method on Mueller-Hinton agar, in accordance with the guidelines of the Clinical and Laboratory Standards Institute (CLSI, 2015). Broth cultures of individual isolates were grown overnight at 37°C, and 200 µl of the standardised broth culture was spread evenly over the surface of Mueller-Hinton agar plates using a sterile glass spreader. Antimicrobial discs were applied to the inoculated surface with approximately 1 cm spacing between discs, and plates were incubated at 37°C for 16-18 hours. Zones of inhibition were measured to the nearest millimetre using a ruler, and isolates were classified as sensitive, intermediate or resistant according to the CLSI zone-diameter interpretive standards for each antimicrobial agent.

**Table 2.** Zone-diameter interpretative standards used for classification of antimicrobial resistance.

| Antimicrobial agent | Symbol | Disc concentration (µg/disc) | Resistant (mm) | Intermediate (mm) | Sensitive (mm) |
| --- | --- | --- | --- | --- | --- |
| Ciprofloxacin | CIP | 5 | ≤ 15 | 16-20 | ≥ 21 |
| Amoxicillin | AMX | 30 | ≤ 13 | 14-17 | ≥ 18 |
| Doxycycline | DO | 30 | ≤ 10 | 11-13 | ≥ 14 |
| Oxytetracycline | TE | 30 | ≤ 11 | 12-14 | ≥ 15 |
| Levofloxacin | LVX | 5 | ≤ 13 | 14-16 | ≥ 17 |
| Neomycin | N | 30 | ≤ 12 | 13-16 | ≥ 17 |
| Erythromycin | E | 15 | ≤ 13 | 14-22 | ≥ 23 |

### 2.7 Maintenance of Stock Culture

Confirmed isolates were preserved in 50% sterile buffered glycerin, prepared by combining equal parts of pure glycerin and phosphate-buffered saline (PBS). A loopful of dense bacterial culture from each isolate was mixed with the buffered glycerin in sterile vials and stored at -20°C for subsequent reference.

## 3. RESULTS

### 3.1 Cultural and Morphological Characteristics

Growth of *E. coli* in nutrient broth was indicated by diffuse turbidity, with pellicle formation observed in a few cases. On nutrient agar, isolates formed smooth, circular, white-to-greyish-white colonies. On EMB agar, colonies were smooth and circular with a characteristic greenish-black colour and metallic sheen, while on MacConkey agar isolates produced smooth, bright pink, lactose-fermenting colonies. On Brilliant Green agar, colonies appeared yellow and the medium changed from red to yellow, and on XLD agar colonies were yellow with the medium changing from yellowish-red to yellow, reflecting partial suppression of xylose, lactose and sucrose degradation. Microscopic examination of Gram-stained smears revealed Gram-negative, short, rod-shaped organisms arranged singly or in pairs, consistent with the typical morphology of *E. coli*.

**Fig. 2.**
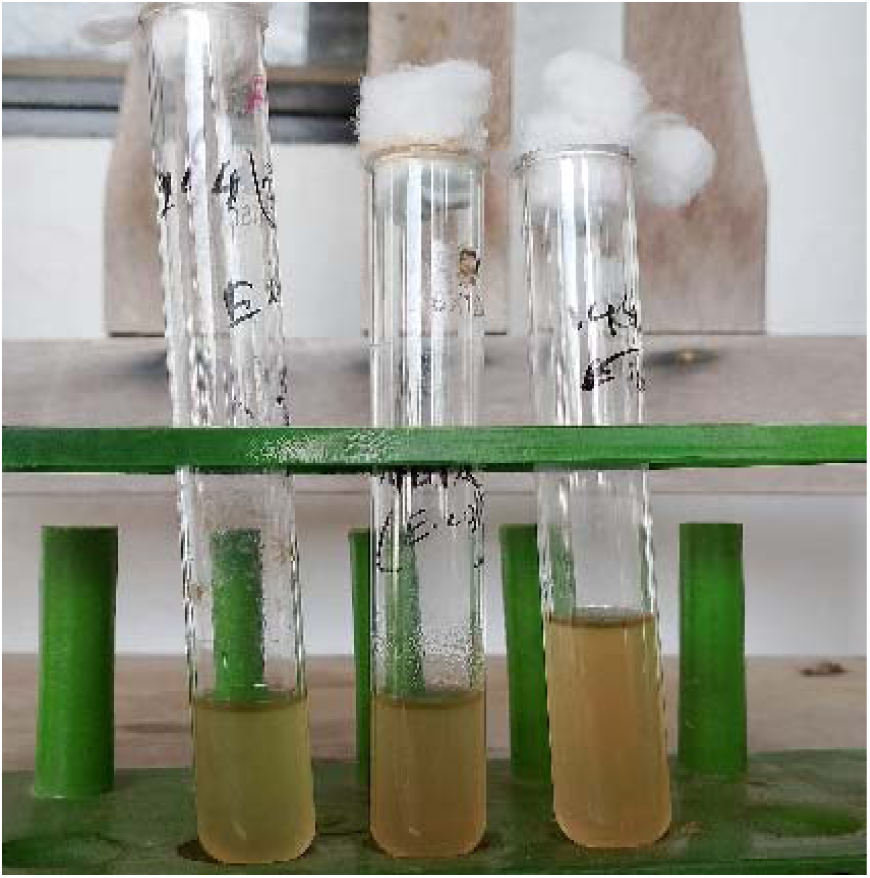
Growth of E. coli in nutrient broth, shown by diffuse turbidity and heavy sediment (left tube: control).

**Fig. 3.**
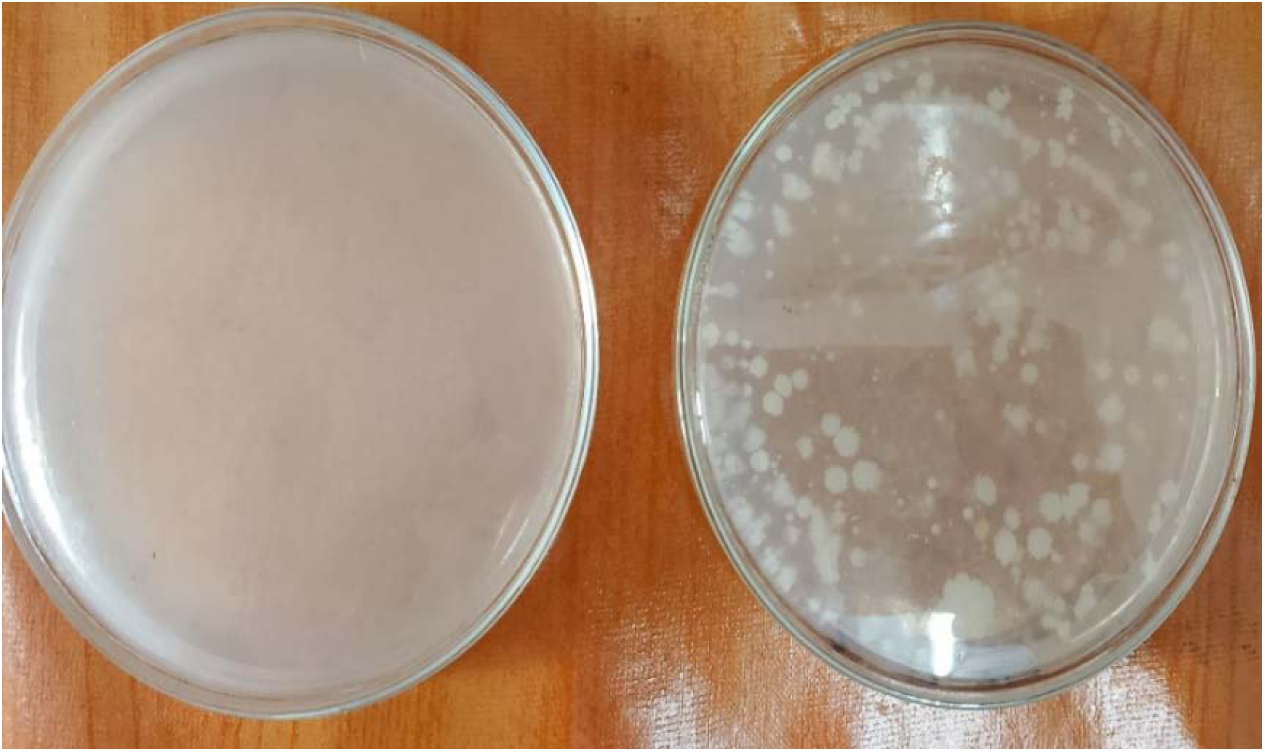
Growth of E. coli on nutrient agar, showing smooth, circular, white to greyish-white colonies (left plate: control).

**Fig. 4.**
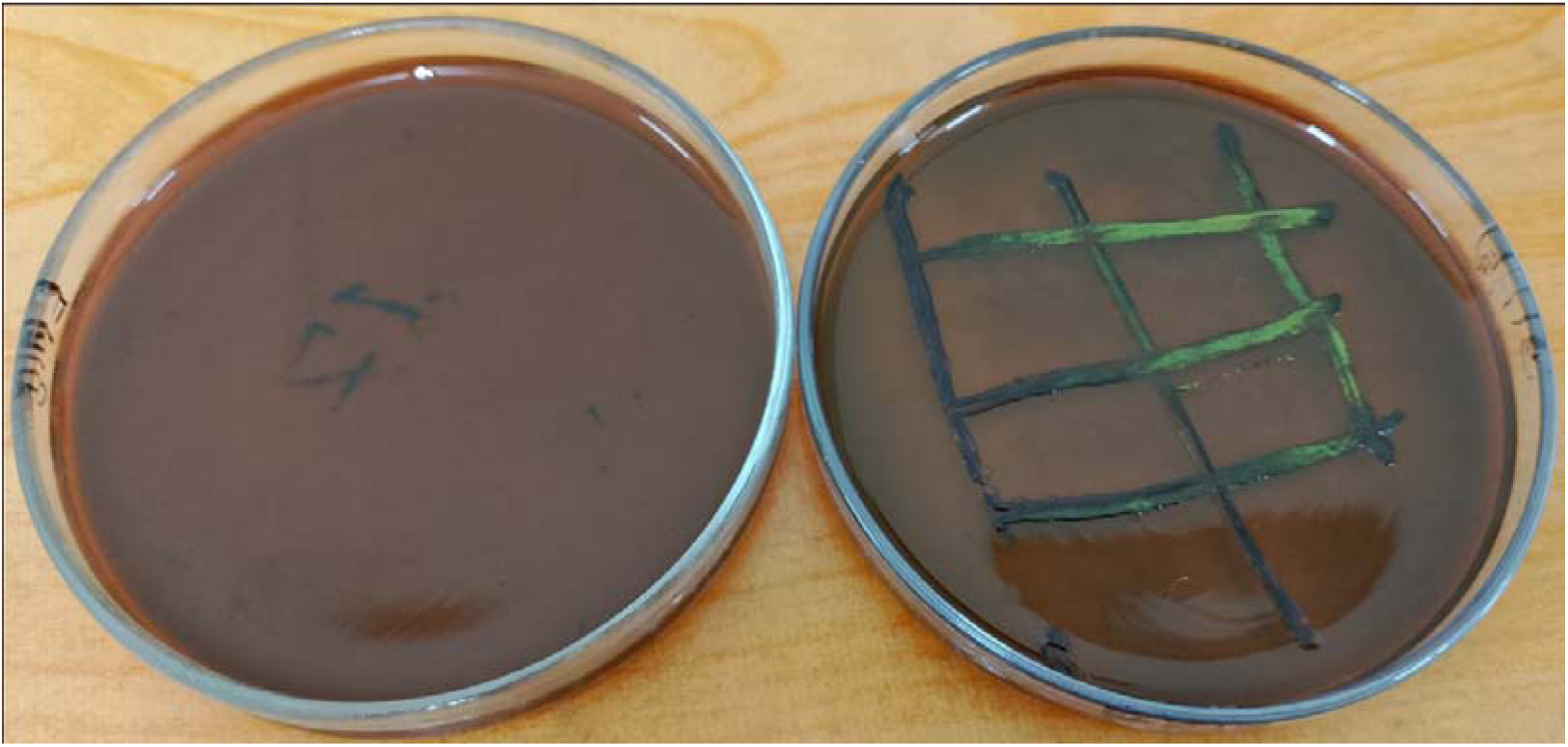
Growth of E. coli on EMB agar, showing greenish-black colonies with characteristic metallic sheen.

**Fig. 5.**
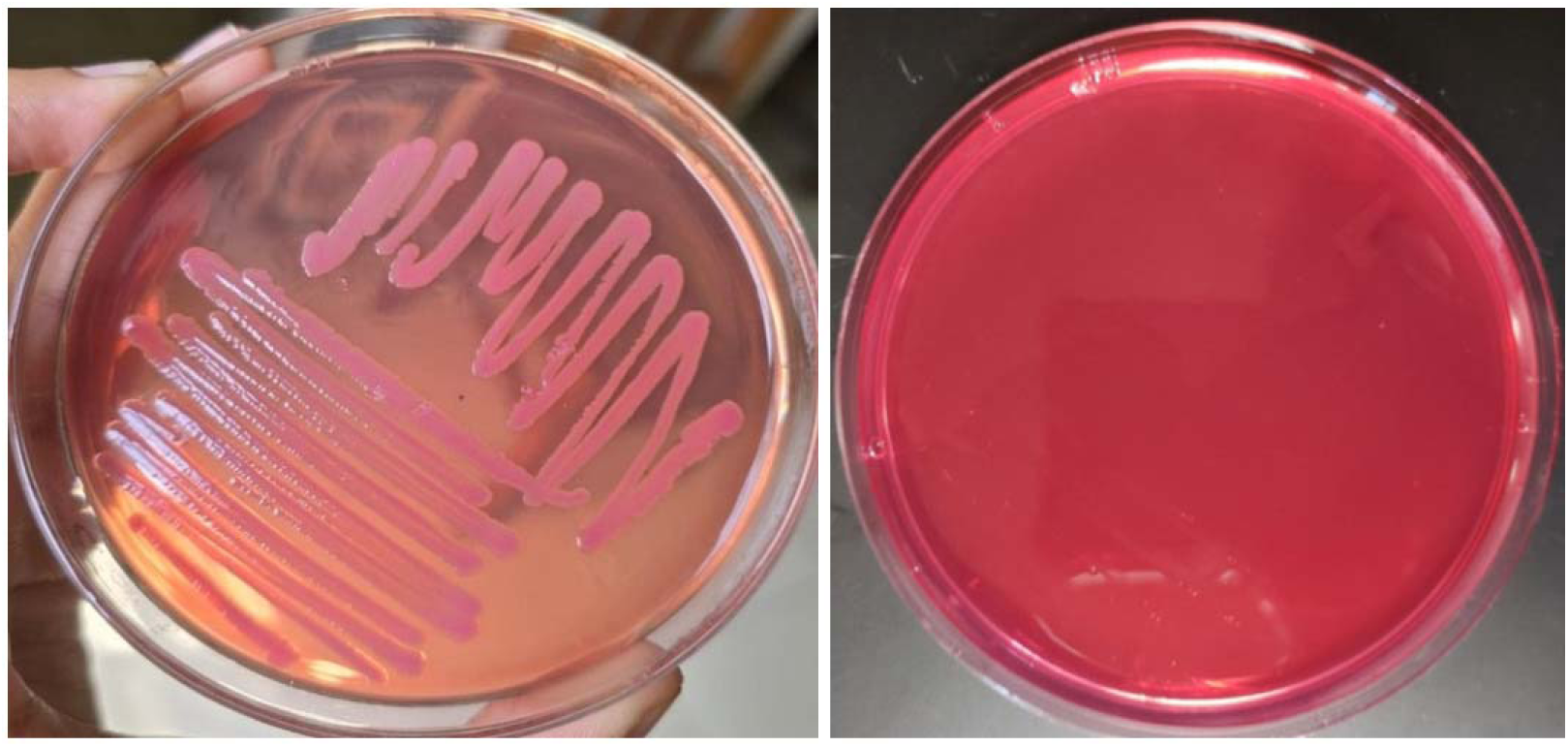
Growth of E. coli on MacConkey agar, showing lactose-fermenting bright pink colonies (right plate: control).

**Fig. 6.**
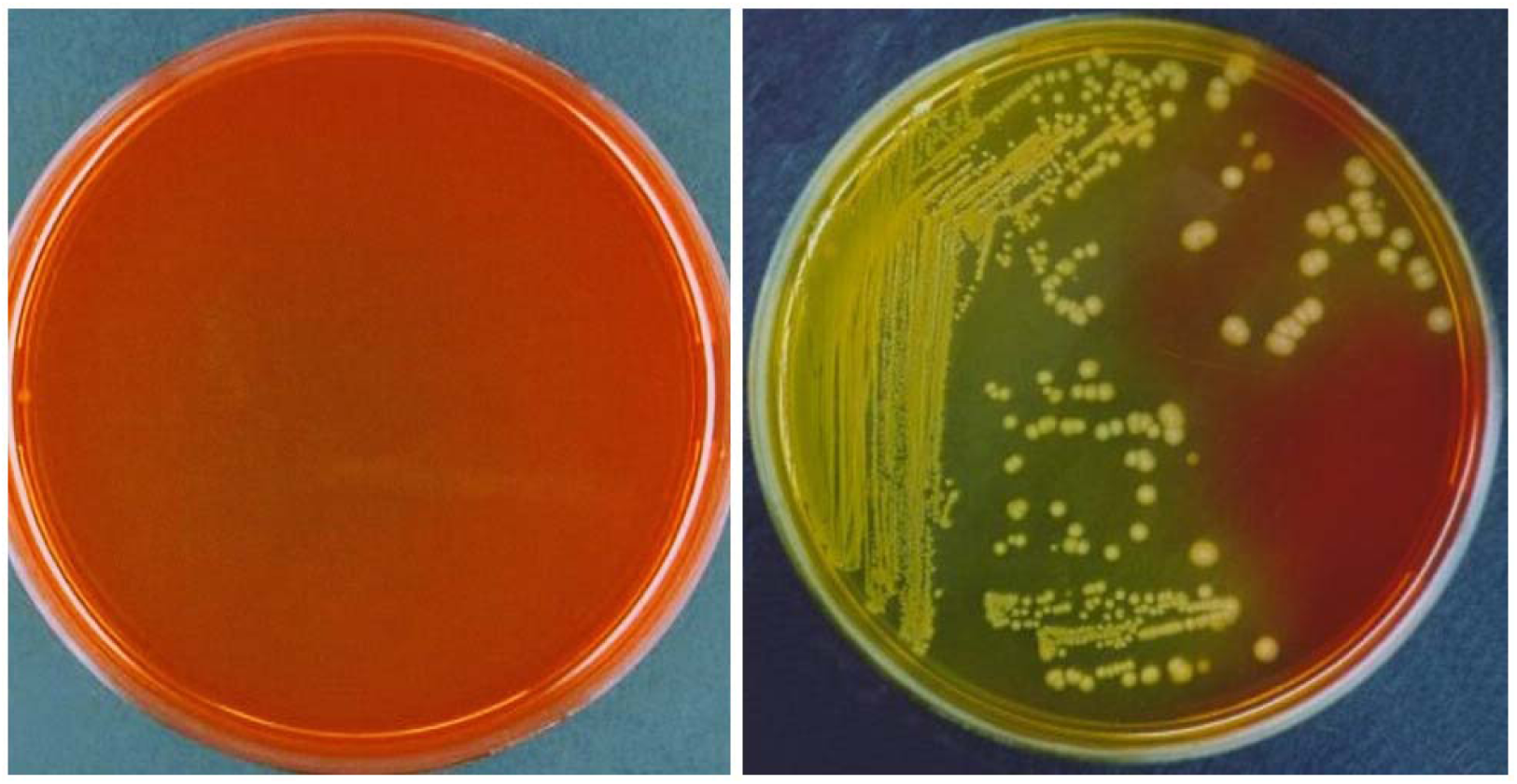
Growth of E. coli on Brilliant Green agar, showing yellow colonies with the medium changing from red to yellow (left plate: control).

**Fig. 7.**
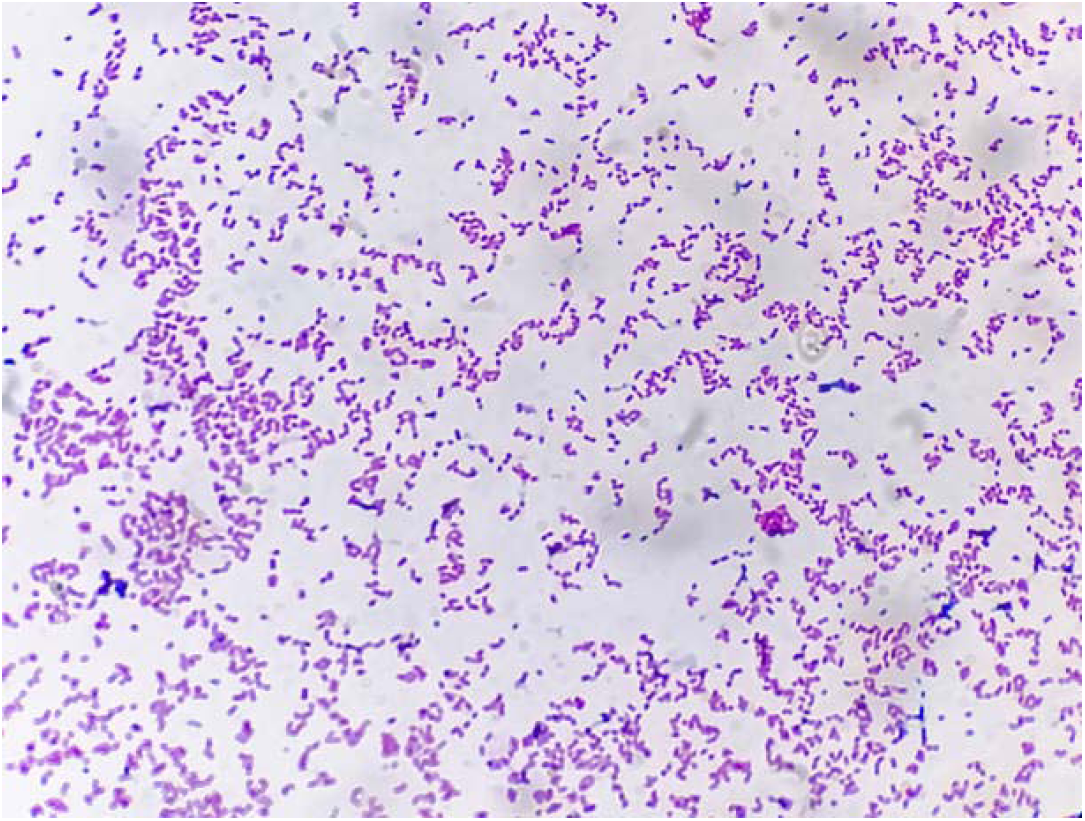
Gram’s staining of E. coli showing Gram-negative, rod-shaped organisms, single or paired, under light microscopy (100x).

**Table 3.** Cultural, staining and motility characteristics of isolated *E. coli* on different media.

| Medium | Colony characteristics |
| --- | --- |
| Nutrient agar | Smooth, circular, white to greyish-white colony |
| EMB agar | Smooth, circular, greenish-black colony with metallic sheen |
| MacConkey agar | Smooth, bright pink coloured colony |
| Brilliant Green agar | Yellow colony; medium colour changed from red to yellow |
| XLD agar | Yellow colony; medium colour changed from yellowish-red to yellow |
| Gram's staining | Gram-negative, short, plump rods, single or paired |
| Motility test | Motile |

### 3.2 Sugar Fermentation and Biochemical Characterization

All isolates fermented the five basic sugars tested-dextrose, maltose, lactose, sucrose and mannitol-with the production of both acid and gas, indicated by a colour change from red to yellow in the medium and gas accumulation in the inverted Durham’s tubes. On biochemical testing, all isolates were catalase-positive and indole-positive, but methyl-red (MR) and Voges-Proskauer (VP) test results, together with Simmons citrate utilisation, were negative. On TSI agar slant, all isolates produced an acidic slant and acidic butt (yellow slant, yellow butt) with gas production, consistent with fermentation of glucose, lactose and sucrose without hydrogen sulphide production.

**Fig. 8.**
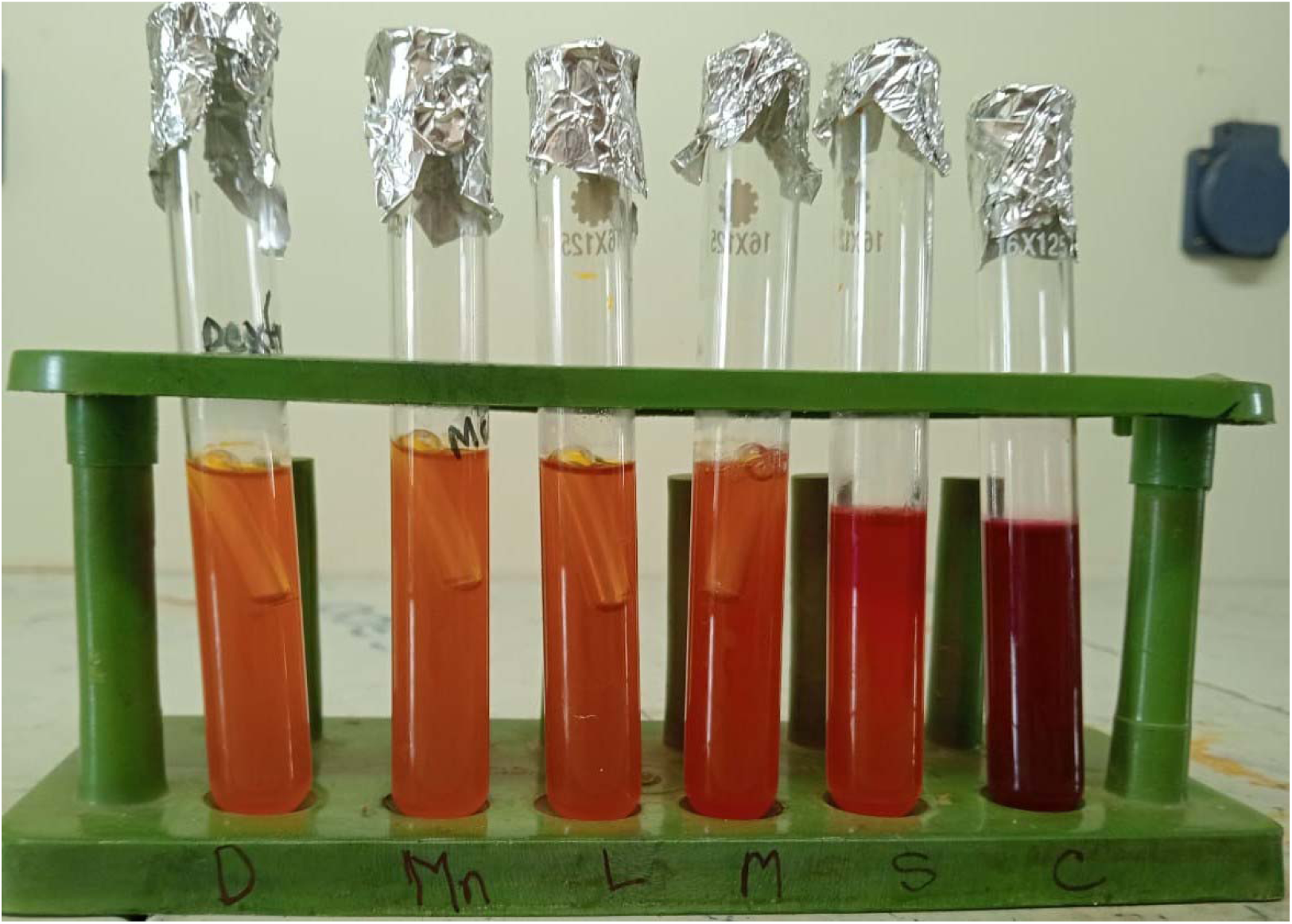
Fermentation of the five basic sugars by E. coli, showing acid and gas production (right tube: control).

**Fig. 9.**
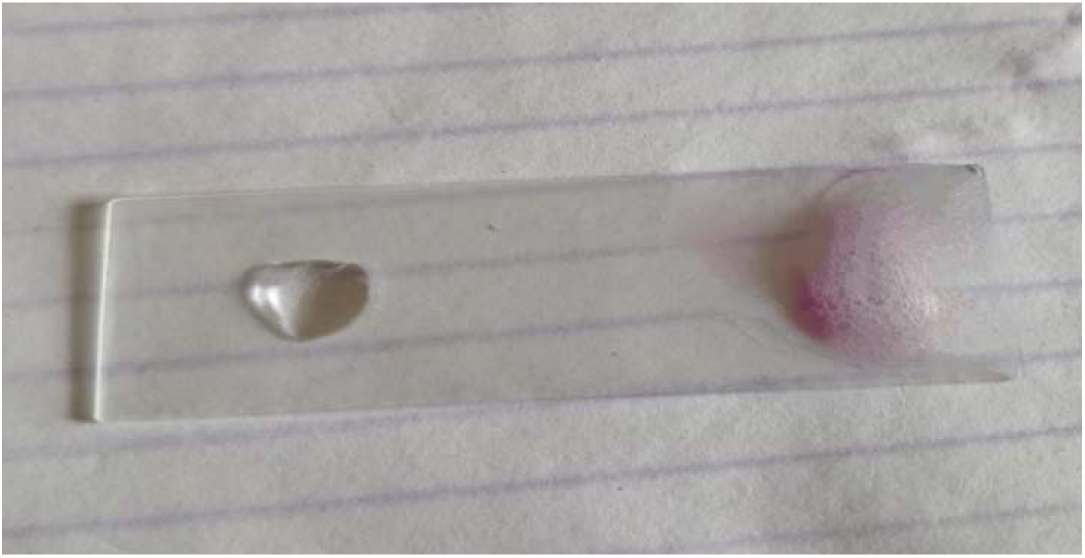
Catalase test showing a positive result indicated by bubble formation (left: control).

**Fig. 10.**
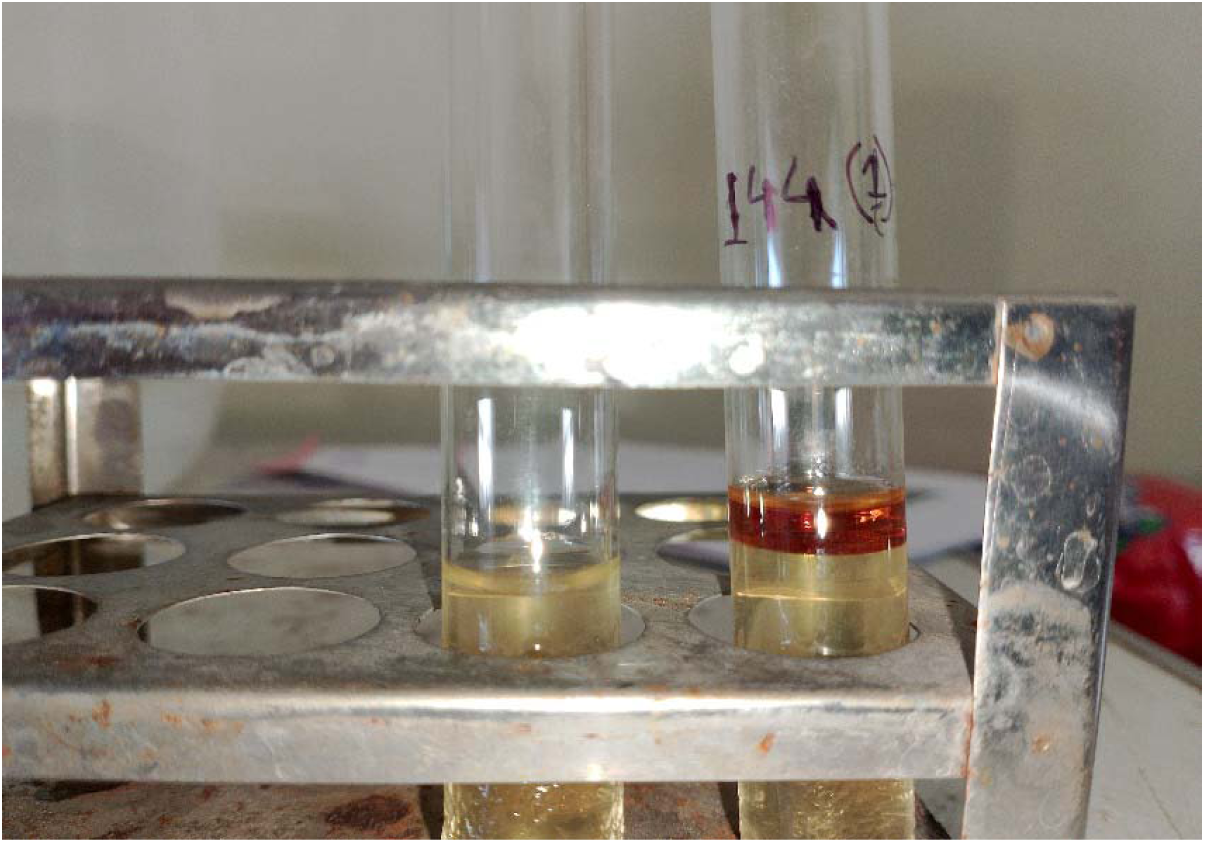
Indole test showing a positive result indicated by a red-violet colour at the surface of the alcohol layer (left: control).

**Fig. 11.**
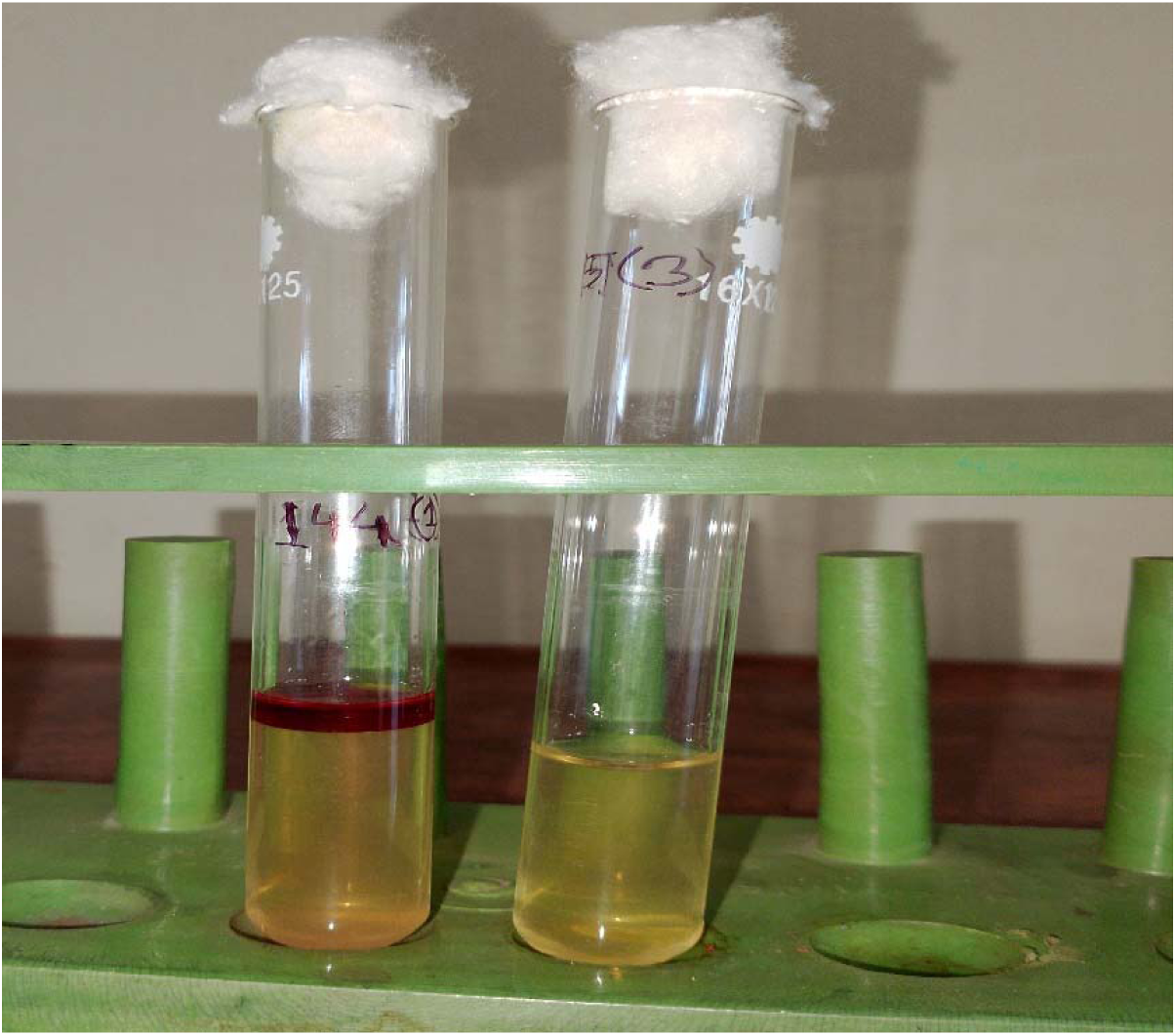
Methyl-red (MR) test showing a positive result indicated by development of a red colour (right: control).

**Fig. 12.**
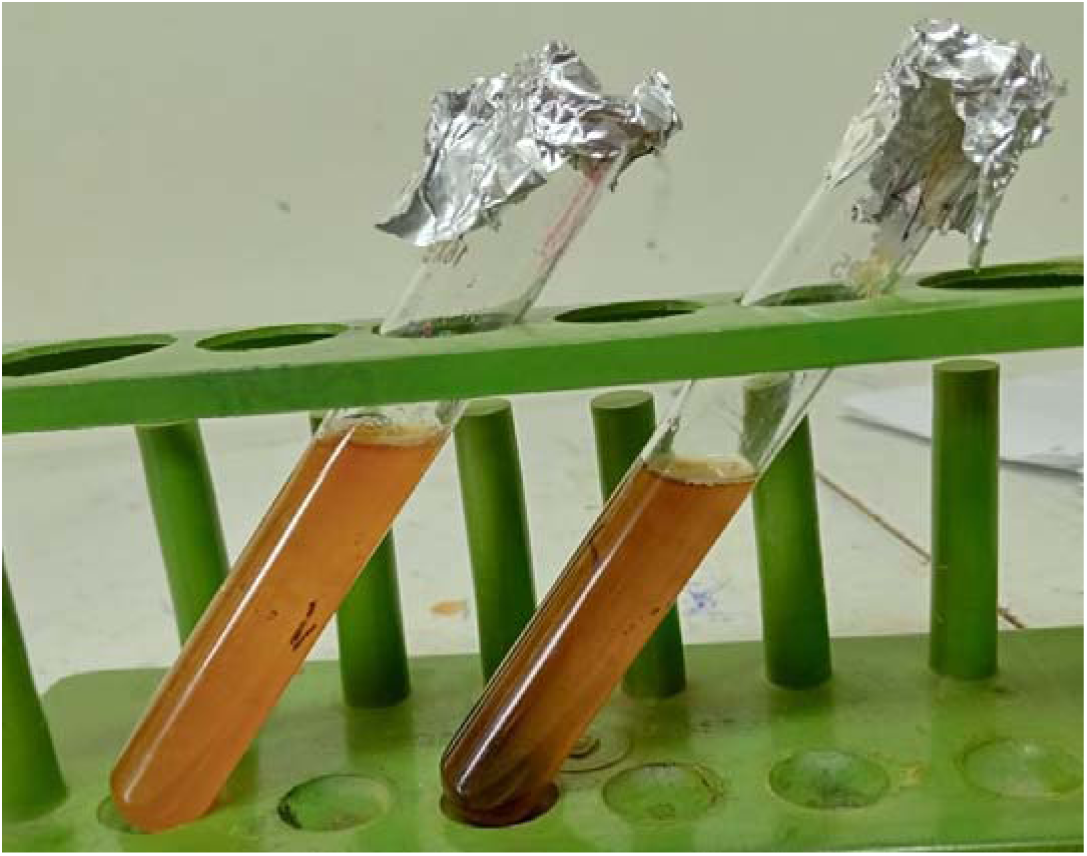
TSI agar slant reaction showing an acidic slant and acidic butt with gas production (left: control).

**Fig. 13.**
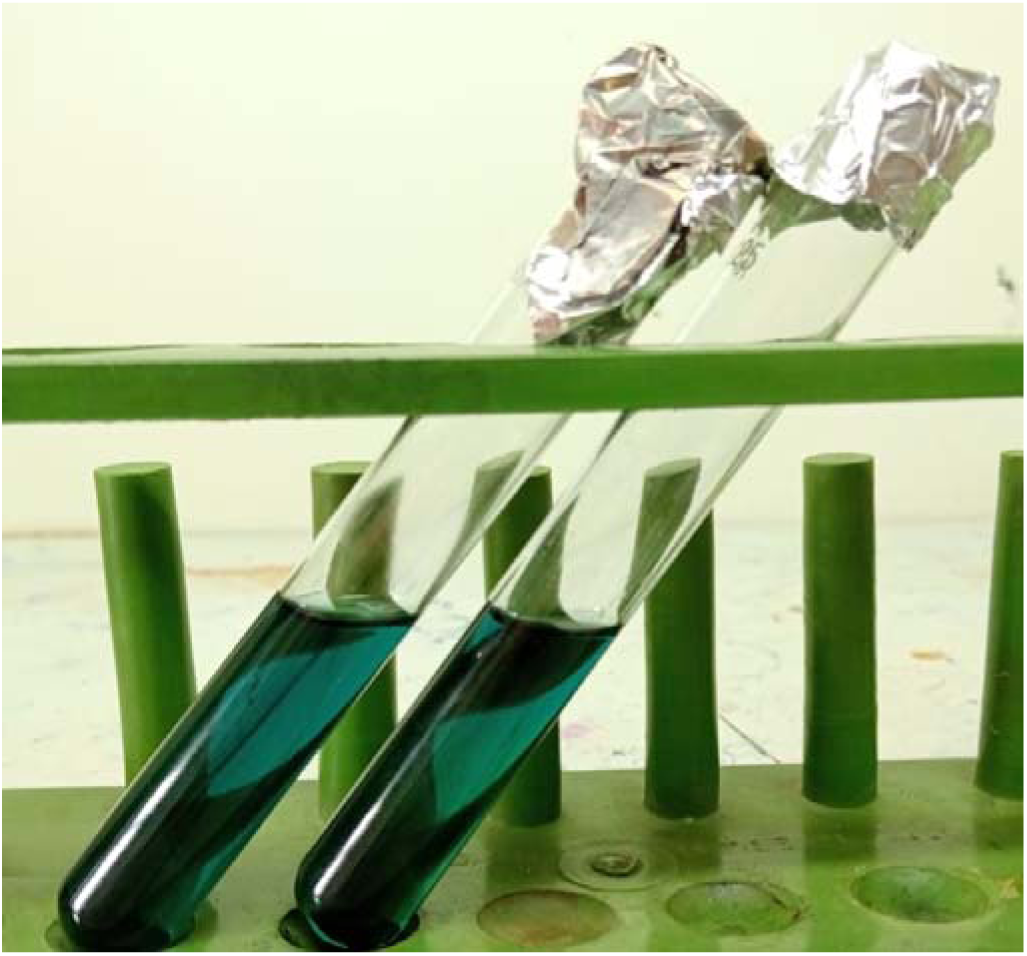
Simmons citrate agar slant reaction showing a negative result (left: control).

**Table 4.**
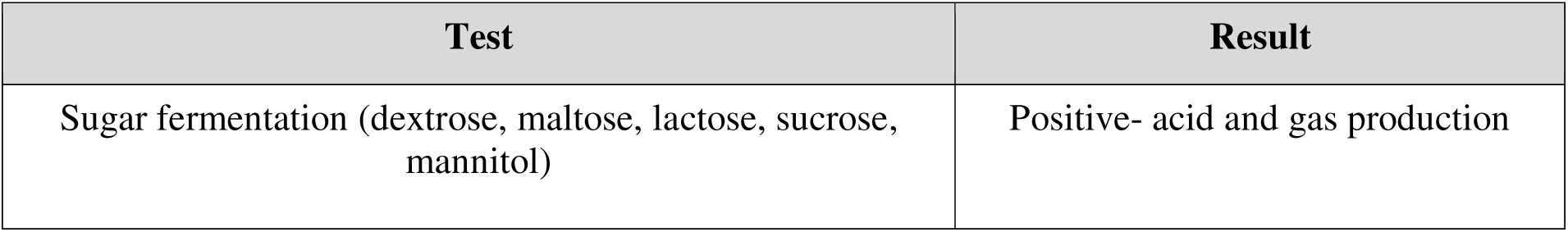
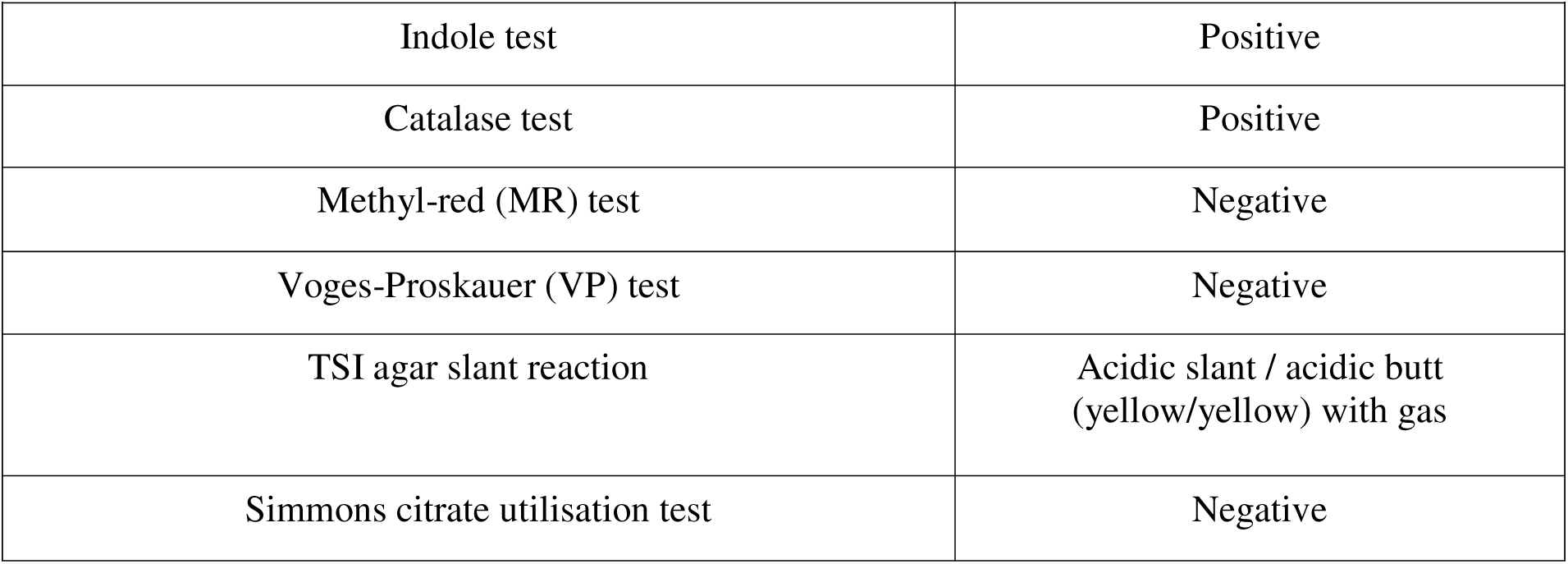
Biochemical properties of isolated *E. coli*.

**Fig. 14.**
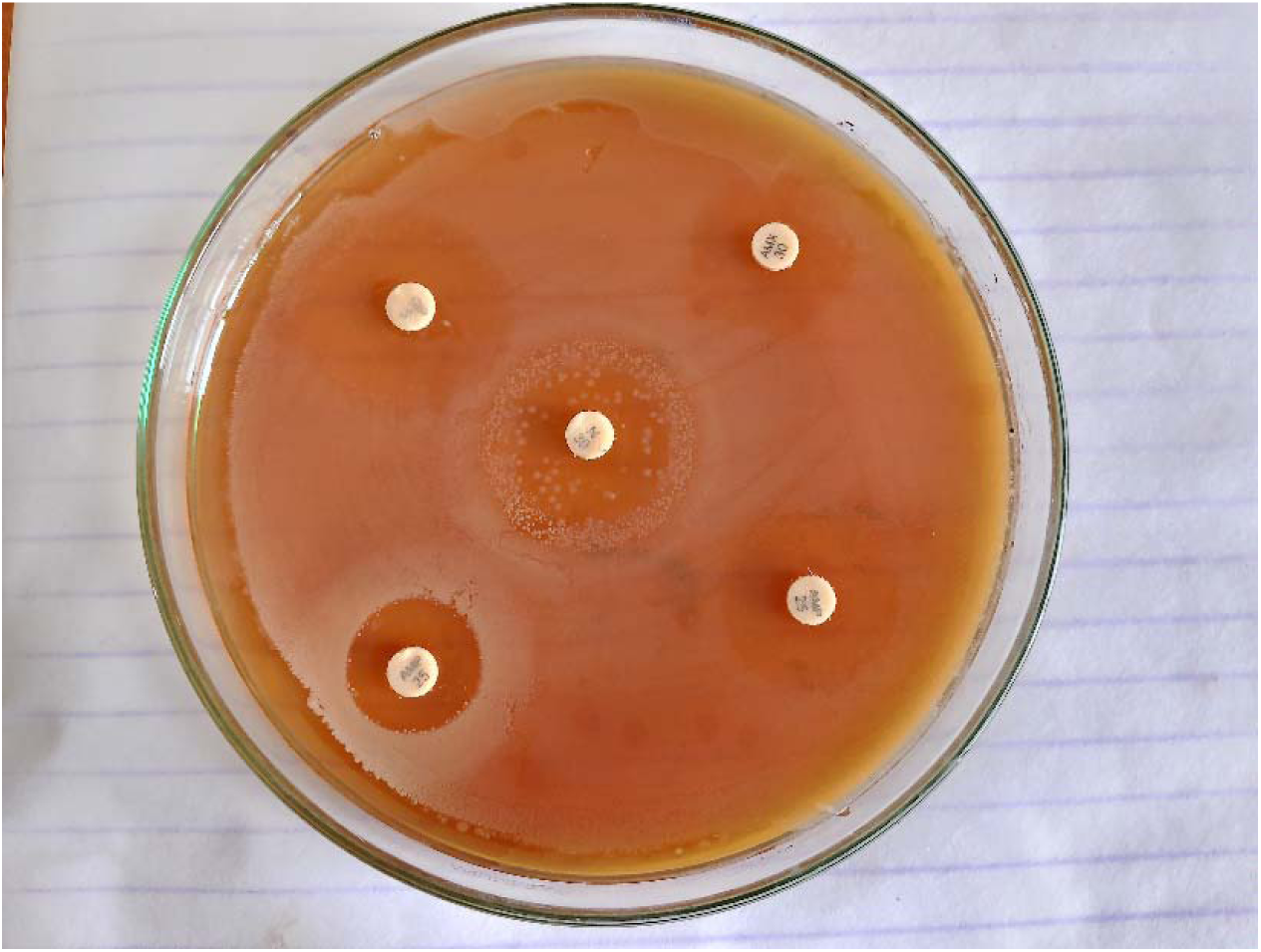
Antibiotic sensitivity and resistance pattern of E. coli on Mueller-Hinton agar, showing resistance to Neomycin and sensitivity to Doxycycline, Ciprofloxacin and Levofloxacin, with intermediate susceptibility to Erythromycin, Amoxicillin and Oxytetracycline.

### 3.3 Overall Prevalence of *E. coli*

Of the 60 environmental samples examined, 33 were positive for *E. coli*, giving an overall prevalence of 55.00% in the live bird market environment of Rajshahi District. Prevalence differed markedly by sample type: 11 of 20 water samples (55.00%), 7 of 20 soil samples (35.00%) and 15 of 20 bird-dropping samples (75.00%) were positive for *E. coli*.

**Table 5.** Overall prevalence of *E. coli* in environmental samples collected from live bird markets of Rajshahi District.

| Sample type | No. of samples examined | No. positive for *E. coli* (%) |
| --- | --- | --- |
| Water | 20 | 11 (55.00%) |
| Soil | 20 | 7 (35.00%) |
| Bird droppings | 20 | 15 (75.00%) |
| Total | 60 | 33 (55.00%) |

**Fig. 15.**
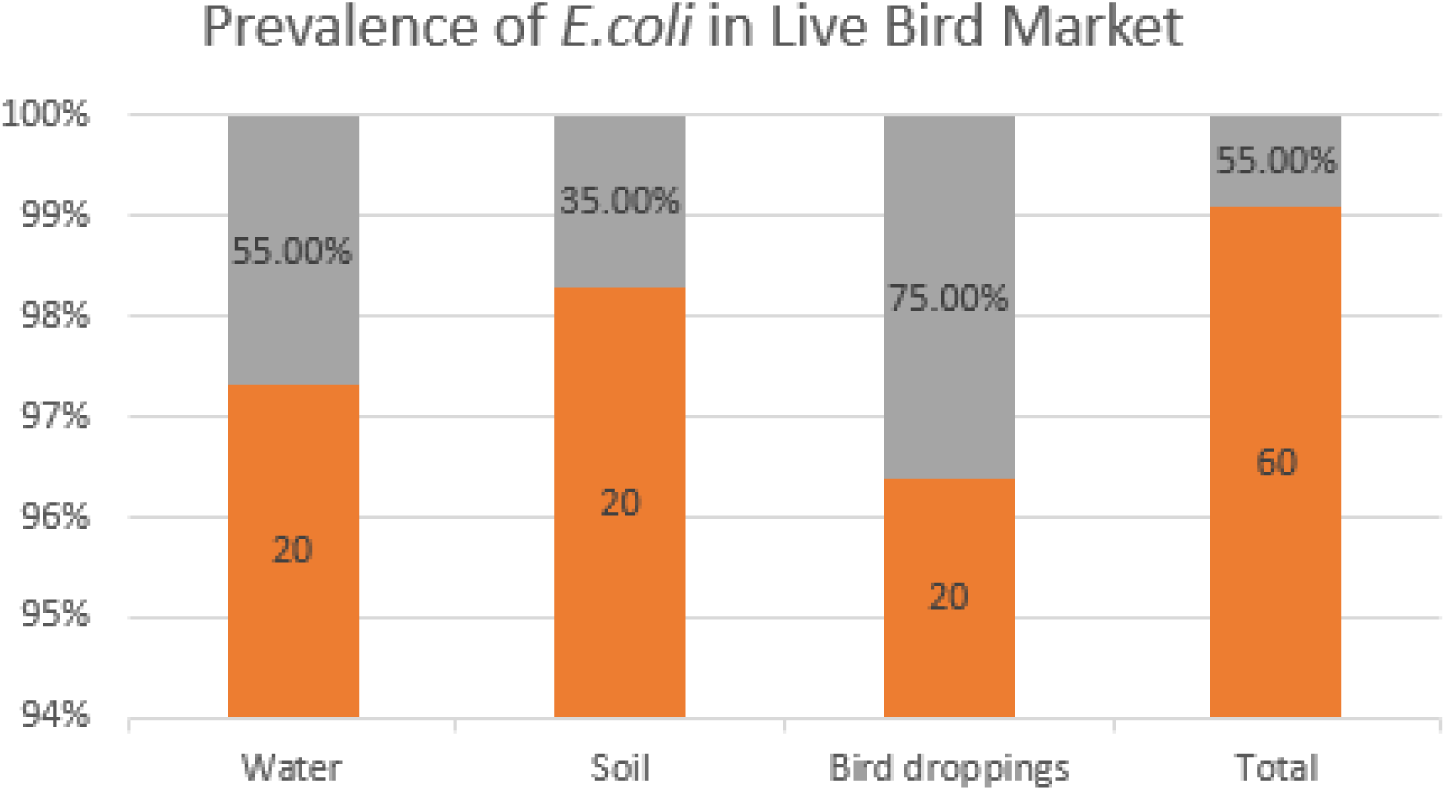
Overall prevalence of E. coli in environmental samples from live bird markets of Rajshahi District.

### 3.4 Antibiogram Profile of Isolated *E. coli*

Antibiotic susceptibility testing of the 33 *E. coli* isolates against seven antimicrobial agents revealed a high degree of resistance to older, broad-spectrum antibiotics and relatively higher sensitivity to newer agents. Resistance was highest to Oxytetracycline (78.79%), followed by Amoxicillin (63.64%), Ciprofloxacin (48.48%), Doxycycline (33.33%), Levofloxacin (9.09%), Erythromycin (9.09%) and Neomycin (6.06%). Correspondingly, sensitivity was highest to Neomycin (60.61%), followed by Levofloxacin (51.51%), Erythromycin (51.51%), Ciprofloxacin (36.36%), Amoxicillin (21.21%), Doxycycline (18.18%) and Oxytetracycline (18.18%). Intermediate sensitivity ranged from 3.03% (Oxytetracycline) to 48.48% (Doxycycline) across the antibiotics tested.

**Table 6.**
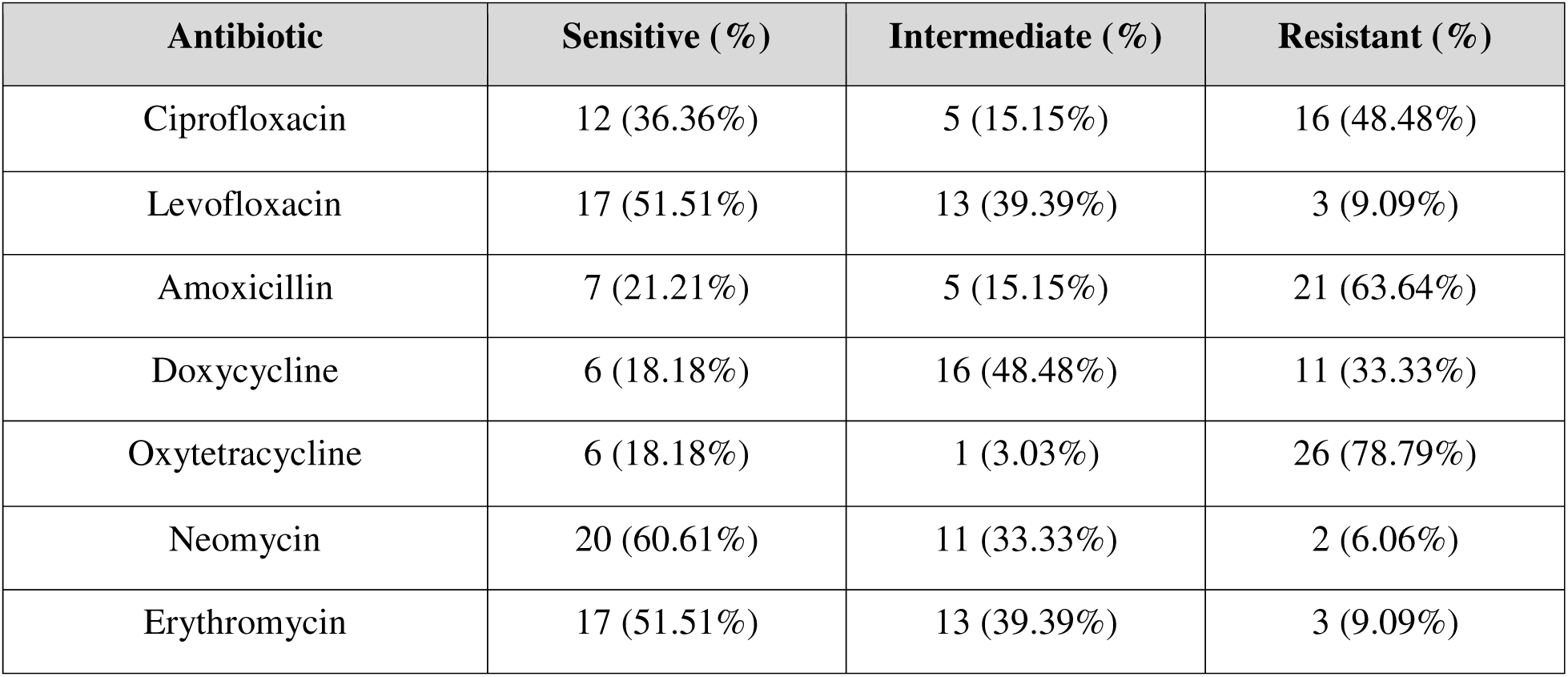
Antibiotic sensitivity and resistance pattern of *E. coli* (n = 33) isolated from th environment of live bird markets.

**Fig. 16.**
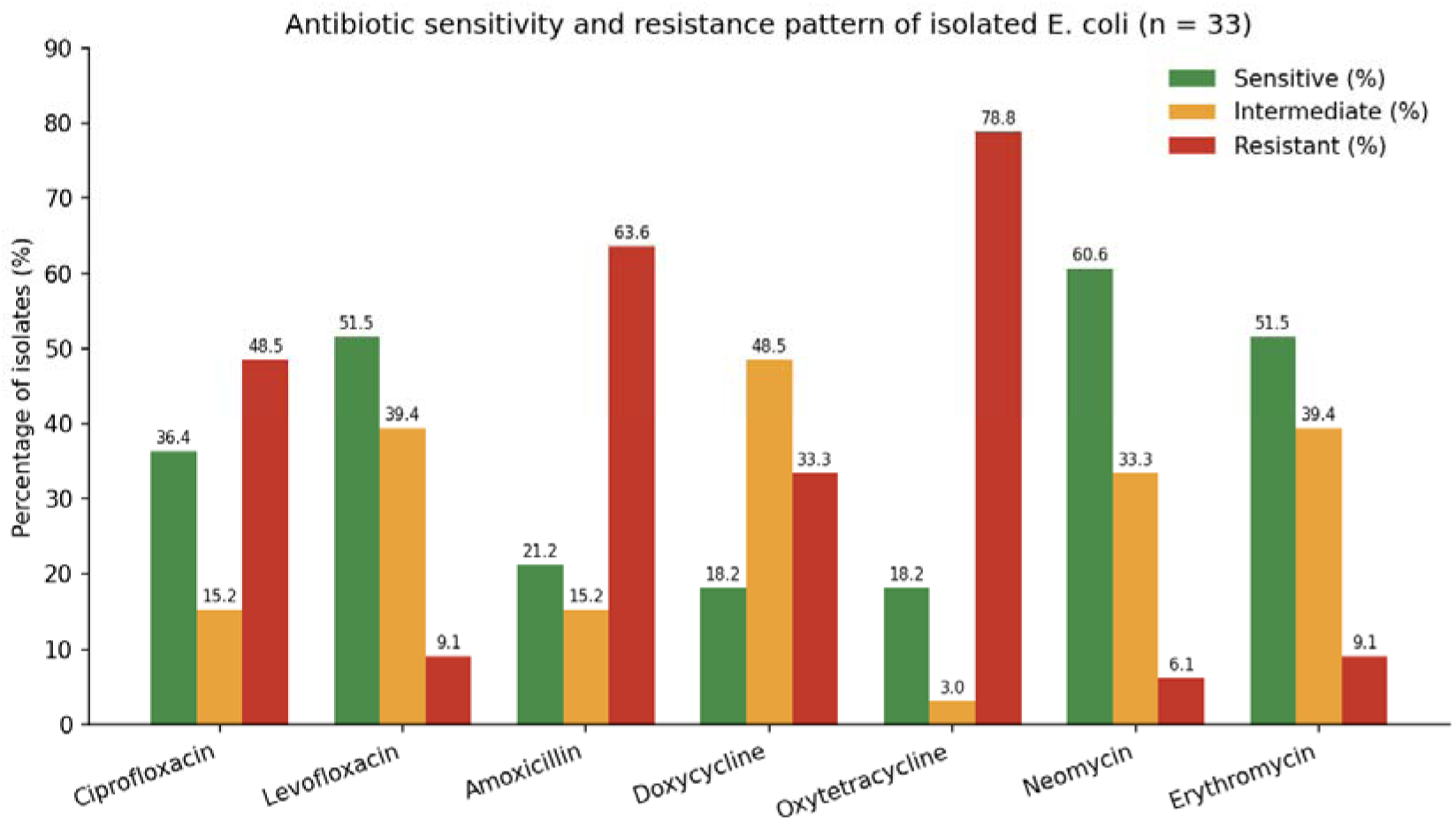
Antibiotic sensitivity and resistance pattern of isolated E. coli (n = 33).

## 4. DISCUSSION

Enterobacteria are a heterogeneous group of Gram-negative rods that naturally colonise the intestinal tracts of birds and other warm-blooded animals. In the present study, isolated *E. coli* displayed the typical short, rod-shaped, Gram-negative morphology, arranged singly or in pairs, and fermented all five basic carbohydrates tested with acid and gas production, consistent with the recognised biochemical profile of the species.

The overall prevalence of *E. coli* recorded in this study (55.00%) is broadly consistent with, though somewhat lower than, several previous reports from live bird markets in Bangladesh and neighbouring countries. Sarker et al. (2019) reported that 37 of 60 samples (61.67%) collected from two live bird markets in Chattogram, Bangladesh, were positive for *E. coli*. Dey et al. (2013) found that 78 of 112 pigeon samples (69.64%) collected from live bird markets in Mymensingh were positive for *E. coli*. Adebowale et al. (2022) reported 56.3% multidrug-resistant *E. coli* among environmental samples from eight live bird markets in Abeokuta, Nigeria, while Surra Gebeyehu et al. (2018) recovered *E. coli* from 32 of 90 pooled samples (58.20%) collected from five live bird markets in Addis Ababa, Ethiopia. Nabil et al. (2020) reported a comparatively lower prevalence of 37.2% among environmental samples from wild-bird live markets on Egypt’s northern coast.

With respect to sample type, the prevalence of *E. coli* recorded in the present study- 55.00% in water, 35.00% in soil and 75.00% in bird-dropping samples- is comparable to findings reported by Blaak et al. (2015), who detected *E. coli* contamination of a poultry farm environment at rates of 81% in rinse water, 60% in dust, 57% in surface water, 55% in soil and 6% in barn air. Da Costa et al. (2008) reported that 22 of 40 wastewater samples (55.7%) from eight poultry slaughterhouses were positive for antibiotic-resistant *E. coli*. Mahmud et al. (2018) recorded a prevalence of 83.08% in cloacal samples from healthy broiler chickens, while Laube et al. (2014) found that 12 of 14 slurry samples (86%) from broiler farms were positive for ESBL/AmpC-producing *E. coli*, with lower positivity in boot swabs (28.8%) and exhaust air samples (7.5%). Parvin et al. (2020) reported *E. coli* in 71.3% of retail chicken meat samples and 79.7% of sewage water samples collected from 64 live bird markets across Bangladesh, while Mamun et al. (2016) found that 49 of 60 cloacal swabs (81.67%) from a wholesale live bird market in Mymensingh were positive for Shigatoxigenic *E. coli*. Dahshan et al. (2015) recorded an *E. coli* prevalence of 79.5% in litter, bird-dropping and water samples collected from broiler farms in Egypt. Taken together, the variation in prevalence reported across these studies most likely reflects differences in sample type, geographic location, season, biosecurity practices and hygiene standards at the point of sampling.

The antibiogram profile obtained in this study, characterised by high resistance to Oxytetracycline (78.79%) and Amoxicillin (63.64%) and comparatively greater sensitivity to Neomycin (60.61%), Levofloxacin (51.51%) and Erythromycin (51.51%), is consistent with the broader pattern of resistance to older, widely used antimicrobials reported elsewhere. Akond et al. (2009) reported that 88%, 82%, 68%, 64%, 58%, 52% and 20% of *E. coli* strains from poultry sources in Bangladesh were resistant to penicillin, ciprofloxacin, erythromycin, ampicillin, tetracycline and chloramphenicol, respectively. Such findings across independent studies of poultry-associated *E. coli* point to widespread, sustained selection pressure from the older tetracycline and beta-lactam classes of antimicrobials, which have historically been the most affordable and most heavily used agents in poultry production.

Variation in the prevalence and resistance patterns of *E. coli* recovered from live bird market environments can plausibly be attributed to differences in biosecurity control, handling practices and sanitary conditions at the farm, transport and market level. Because birds and their environmental substrates in live bird markets are subject to continual mixing from multiple source flocks, both vertical transmission from infected parent stock and horizontal contamination within the highly trafficked market environment, where iron availability and organic load favour rapid bacterial proliferation, are likely to contribute to the overall contamination burden. Even though the enteric bacteria isolated from these environmental sources are not, in themselves, invariably pathogenic, consumers who handle or consume raw or undercooked poultry products originating from contaminated markets nonetheless remain at risk of exposure to multidrug-resistant organisms and their associated resistance determinants.

## 5. CONCLUSION

This study demonstrated a substantial prevalence of Escherichia coli (55.00%) in the environmental samples of live bird markets across Rajshahi District, with the highest contamination detected in bird-dropping samples, followed by water and soil. The isolates showed considerable resistance to oxytetracycline, amoxicillin, ciprofloxacin and doxycycline, while comparatively higher susceptibility was observed to neomycin, levofloxacin and erythromycin. These findings highlight the potential role of live bird market environments as reservoirs of E. coli and antimicrobial-resistant bacteria. Strengthening sanitation, environmental hygiene, biosecurity and hygienic poultry-handling practices, together with prudent and judicious antimicrobial use, is essential to reduce the spread of antimicrobial-resistant E. coli and associated public health risks. Further studies incorporating molecular characterization of antimicrobial resistance and virulence determinants are recommended to better assess the epidemiological and public health significance of these isolates.

## Acknowledgements

The authors gratefully acknowledge the Department of Veterinary & Animal Sciences, University of Rajshahi, Bangladesh, for providing laboratory facilities and technical support throughout this study. The authors also sincerely thank the concerned live bird market authorities, market personnel, and other relevant stakeholders in Rajshahi District for their cooperation and assistance during sample collection.

## Ethical Approval

Ethical approval for this study was obtained from the Institutional Animal, Medical Ethics, Biosafety and Biosecurity Committee (IAMEBBC), University of Rajshahi, Bangladesh, before commencement of the study. The study involved the collection and microbiological examination of environmental samples, including water, soil, and bird-dropping samples, from live bird markets. No live animals were experimentally handled, restrained, or subjected to invasive procedures during the study.

## Informed Consent

Permission was obtained from the respective live bird market authorities or responsible personnel prior to environmental sample collection. No personal information from market personnel or other individuals was collected or reported in this study.

## Data Availability

The data supporting the findings of this study are available from the corresponding author upon reasonable request. All data generated or analyzed during this study are included in this article or can be obtained from the corresponding author upon reasonable request.

## Author Contributions

Mst. Nahida Akter: Conceptualization, Literature review, Sample collection, Data curation, Laboratory experiments, Data analysis, Methodology, Statistical analysis, Writing-original draft.

Md. Rimon Bhuiyan: Investigation, Data curation, Data analysis, Laboratory experiments, Literature review, Visualization, Writing-original draft, Writing-review and editing.

Md. Sohel Rana, Rubina Khatun: Sample collection, Data curation, Laboratory experiments, Investigation, Writing-original draft.

Anna Purnna Ray: Data curation, Validation, Investigation.

K.M. Mozaffor Hossain: Conceptualization, Methodology, Supervision, Project administration, Writing-review and editing, Final approval of the manuscript.

## REFERENCES

Adebowale, O., Makanjuola, M., Bankole, N., Olasoju, M., Alamu, A., Kperegbeyi, E. and Fasina, F.O. (2022) Multi-drug resistant *Escherichia coli*, biosecurity and anti-microbial use in live bird markets, Abeokuta, Nigeria. Antibiotics, 11(2), p.253.

Akond, M.A., Alam, S., Hassan, S.M.R. and Shirin, M. (2009) Antibiotic resistance of *Escherichia coli* isolated from poultry and poultry environment of Bangladesh. Internet Journal of Food Safety, 11, pp.19–23.

Alexander, D.J., Younus, M., Maqbool, A., Khan, I. and Umar, S. (2017) Pathological alterations during co-infection of Newcastle disease virus with *Escherichia coli* in broiler chicken. Pakistan Journal of Zoology, 49(6), p1953

Amara, A., Ziani, Z. and Bouzoubaa, K. (1995) Antibioresistance of *Escherichia coli* strains isolated in Morocco from chickens with colibacillosis. Veterinary Microbiology, 43(4), pp.325–330.

Armstrong, G. L., Hollingsworth, J., & Morris Jr, J. G. (1996). Emerging foodborne pathogens: Escherichia coil O157: H7 as a model of entry of a new pathogen into the food supply of the developed world. Epidemiologic reviews, 18(1), 29–51.

Baergen, R. (2013) Placental Pathology: An Issue of Surgical Pathology Clinics, Vol. 6, No. 1.Philadelphia: Elsevier Health Sciences.

Balière, C., Rincé, A., Blanco, J., Dahbi, G., Harel, J., Vogeleer, P. and Gourmelon, M. (2015) Prevalence and characterization of Shiga toxin-producing and enteropathogenic *Escherichia coli* in shellfish-harvesting areas and their watersheds. Frontiers in Microbiology, 6, p.1356.

Blaak, H., van Hoek, A.H., Hamidjaja, R.A., van der Plaats, R.Q., Kerkhof-de Heer, L., de Roda Husman, A.M. and Schets, F.M. (2015) Distribution, numbers, and diversity of ESBL-producing *E. coli* in the poultry farm environment. PLOS ONE, 10(8), p.e0135402.

Cheesbrough, M. (1985) Microbiology, in Medical Laboratory Manual for Tropical Countries, Vol. 2. London: English Language Book Society, pp.225–392.

CLSI (2015) Performance Standards for Antimicrobial Susceptibility Testing. Wayne, PA: Clinical and Laboratory Standards Institute.

Da Costa, P.M., Vaz-Pires, P. and Bernardo, F. (2008) Antimicrobial resistance in *Escherichia coli* isolated in wastewater and sludge from poultry slaughterhouse wastewater plants. Journal of Environmental Health, 70(7), pp.40–45.

Dahshan, H., Abd-Elall, A.M.M., Megahed, A.M., Abd-El-Kader, M.A. and Nabawy, E.E. (2015) Veterinary antibiotic resistance, residues, and ecological risks in environmental samples obtained from poultry farms, Egypt. Environmental Monitoring and Assessment, 187, pp.1–10.

Daini, O.A., Ogbolu, O.D. and Ogunledun, A. (2005) Quinolones resistance and R-plasmids of some gram negative enteric bacilli. African Journal of Clinical and Experimental Microbiology, 6(1), pp.14–20.

Davies, J. (1994) Inactivation of antibiotics and the dissemination of resistance genes. Science, 264(5157), pp.375–382.

Dey, R.K., Khatun, M.M., Islam, M.A. and Hosain, M.S. (2013) Prevalence of multidrug resistant *Escherichia coli* in pigeon in Mymensingh, Bangladesh. Microbes and Health, 2(1), pp.5–7.

DLS (2021) Demand, Production and Availability of Milk, Meat and Eggs, 2020-2021. Dhaka: Department of Livestock Services.

Ferri, M., Ranucci, E., Romagnoli, P. and Giaccone, V. (2017) Antimicrobial resistance: a global emerging threat to public health systems. Critical Reviews in Food Science and Nutrition, 57(13), pp.2857–2876.

Harwood, V. J., Staley, C., Badgley, B. D., Borges, K., & Korajkic, A. (2014). Microbial source tracking markers for detection of fecal contamination in environmental waters: relationships between pathogens and human health outcomes. FEMS microbiology reviews, 38(1), 1–40.

Hasina, B. (2006) Enteropathotypic Characterization of *Escherichia coli* Isolated from Diarrhoeic Calves and their Antibiogram Study. MS thesis, Department of Microbiology and Hygiene, Bangladesh Agricultural University, Mymensingh.

Hassan, M.M., Amin, K.B., Ahaduzzaman, M., Alam, M., Faruk, M.S. and Uddin, I. (2014) Antimicrobial resistance pattern against *E. coli* and Salmonella in layer poultry. Research Journal of Veterinary Practitioners, 2(2), pp.30–35.

Islam, M., Sabrin, M.S., Kabir, M.H.B. and Aftabuzzaman, M.D. (2018) Antibiotic sensitivity and resistant pattern of bacteria isolated from table eggs of commercial layers considering food safety issue. Asian Journal of Medical and Biological Research, 4(4), pp.323–329.

Jang, J., Hur, H.G., Sadowsky, M.J., Byappanahalli, M.N., Yan, T. and Ishii, S. (2017) Environmental *Escherichia coli*: ecology and public health implications - a review. Journal of Applied Microbiology, 123(3), pp.570–581.

Kaper, J.B., Nataro, J.P. and Mobley, H.L. (2004) Pathogenic *Escherichia coli*. Nature Reviews Microbiology, 2(2), pp.123–140.

Klein, E.Y., Van Boeckel, T.P., Martinez, E.M., Pant, S., Gandra, S., Levin, S.A. and Laxminarayan, R. (2018) Global increase and geographic convergence in antibiotic consumption between 2000 and 2015. Proceedings of the National Academy of Sciences, 115(15), pp.E3463–E3470.

Kolář, M., Urbánek, K. and Látal, T. (2001) Antibiotic selective pressure and development of bacterial resistance. International Journal of Antimicrobial Agents, 17(5), pp.357–363.

Laube, H., Friese, A., Von Salviati, C., Guerra, B. and Rösler, U. (2014) Transmission of ESBL/AmpC-producing *Escherichia coli* from broiler chicken farms to surrounding areas. Veterinary Microbiology, 172(3-4), pp.519–527.

Looft, T. and Allen, H.K. (2012) Collateral effects of antibiotics on mammalian gut microbiomes. Gut Microbes, 3(5), pp.463–467.

Mahmud, S., Nazir, K.N.H. and Rahman, M.T. (2018) Prevalence and molecular detection of fluoroquinolone-resistant genes (qnrA and qnrS) in *Escherichia coli* isolated from healthy broiler chickens. Veterinary World, 11(12), p.1720.

Mamun, M.M., Parvej, M.S., Ahamed, S., Hassan, J., Nazir, K.H.M.N.H., Nishikawa, Y. and Rahman, M.T. (2016) Prevalence and characterization of shigatoxigenic *Escherichia coli* in broiler birds in Mymensingh. Bangladesh Journal of Veterinary Medicine, 14(1), pp.5–8.

Nabil, N.M., Erfan, A.M., Tawakol, M.M., Haggag, N.M., Naguib, M.M. and Samy, A. (2020) Wild birds in live bird markets: potential reservoirs of enzootic avian influenza viruses and antimicrobial resistant Enterobacteriaceae in northern Egypt. Pathogens, 9(3), p.196.

Neu, H.C. (1992) The crisis in antibiotic resistance. Science, 257(5073), pp.1064–1073.

Osterblad, M., Hakanen, A., Manninen, R., Leistevuo, T., Peltonen, R., Meurman, O. and Kotilainen, P. (2000) A between-species comparison of antimicrobial resistance in enterobacteria in fecal flora. Antimicrobial Agents and Chemotherapy, 44(6), pp.1479–1484.

Parvin, M.S., Talukder, S., Ali, M.Y., Hasan, M.M., Rahman, M.T. and Islam, M.T. (2020) Exploring multidrug resistant *E. coli* carrying extended-spectrum β-lactamase from retail chicken meat and live bird market sewage in Bangladesh. International Journal of Infectious Diseases, 101, p.23.

Read, A.F. and Woods, R.J. (2014) Antibiotic resistance management. Evolution, Medicine, and Public Health, 2014(1), p.147.

Salyers, A.A., Gupta, A. and Wang, Y. (2004) Human intestinal bacteria as reservoirs for antibiotic resistance genes. Trends in Microbiology, 12(9), pp.412–416.

Sarker, M.S., Mannan, M.S., Ali, M.Y., Bayzid, M., Ahad, A. and Bupasha, Z.B. (2019) Antibiotic resistance of *Escherichia coli* isolated from broilers sold at live bird markets in Chattogram, Bangladesh. Journal of Advanced Veterinary and Animal Research, 6(3), p.272.

Surra Gebeyehu, S.G., Dereje Tulu, D.T. and Chaluma Negera, C.N. (2018) Isolation and identification of *Escherichia coli*, *Salmonella* sp. and *Pasteurella* sp. from holding grounds of live-bird markets at Addis Ababa, Ethiopia. African Journal of Microbiology Research, 12(31), pp.754–760.

Tenaillon, O., Skurnik, D., Picard, B. and Denamur, E. (2010) The population genetics of commensal *Escherichia coli*. Nature Reviews Microbiology, 8(3), pp.207–217.

WHO (2020) Antimicrobial Resistance. Geneva: World Health Organization.

